# Nearest neighbor search on embeddings rapidly identifies distant protein relations

**DOI:** 10.1101/2022.09.04.506527

**Authors:** Konstantin Schütze, Michael Heinzinger, Martin Steinegger, Burkhard Rost

## Abstract

Since 1992, all state-of-the-art (SOTA) methods for fast and sensitive identification of evolutionary, structural, and functional relations between proteins (also referred to as “homology detection”) use sequences and sequence-profiles (PSSMs). Protein Language Models (pLMs) generalize sequences, possibly capturing the same constraints as PSSMs, e.g., through embeddings. Here, we explored how to use such embeddings for nearest neighbor searches to identify relations between protein pairs with diverged sequences (remote homology detection for levels of <20% pairwise sequence identity, PIDE). While this approach excelled for proteins with single domains, we demonstrated the current challenges applying this to multi-domain proteins and presented some ideas how to overcome existing limitations, in principle. We observed that sufficiently challenging data set separations were crucial to provide deeply relevant insights into the behavior of nearest neighbor search when applied to the protein embedding space, and made all our methods readily available for others.

## 1 Introduction

### Homology detection

Any investigation of a query protein, Q, beginning with its sequence starts by the identification of evolutionary, structural, and functional relations between Q and all other proteins for which relevant experimental annotations exist. This investigation is often, and slightly misleadingly, labelled as “*homology detection*” inspired by the terminology introduced to describe the analogy of organs between different species (Owen, 1848). *Homology* as a term describing the similarity between proteins typically inherits the concept of “related by evolution” from the original comparison of species (Darwin, 1859). In practice, “related by evolution” is often replaced by “similar (or identical) structure (or function)” as “structural similarity” is less ambiguous to quantify than “evolutionary relation”. This ambivalence also pertains to our view of “protein space”: maps showing relations between proteins are specific to particular definitions, e.g., the map of protein structure domains, or of functional units, or of evolutionary relations. The reproducibility of such maps increases with quantifiability, and is highest for protein structure because the knowledge of structure enables parsing proteins into their compact, independently foldable constituents, namely structural domains. Searches for relations in protein space root almost any drug development and are crucial for the success of the breakthrough prediction of protein structure prediction by *AlphaFold2* (Jumper et al., 2021; Mirdita et al., 2021; Marx, 2022).

### From sequence to feature similarity

With growing databases, speed has become THE major challenge for methods detecting *homology* by comparing the sequence of a query protein Q to all sequences in a database DB. All successful fast solutions from the classic *BLAST*/*PSI-BLAST*, over to *MMseqs2* and *Diamond2*, (Altschul et al., 1997; Steinegger and Söding, 2017; Buchfink et al., 2021) essentially follow three steps. (1) Fast: Initialize search using sequence fragments with typically 3-10 consecutive residues (k-mer), i.e., by finding database sequences with k-mers identical or near-identical to the query. (2) Slower: Expand short k-mer matches (*hits*) through un-gapped alignment. (3) Slowest: Refine alignment for subset of Q-DB pairs with un-gapped scores above a predefined threshold (Step #2) through resource-intensive Smith-Waterman (Smith et al., 1981). Ultimately, the first two steps pre-filter the finding of homologs, while the third generates the actual alignment yielding an E-value for each hit. This E-value estimates how many hits with identical are expected by chance (Karlin and Altschul, 1990). Following others, *MMseqs2* ((Steinegger and Söding, 2017), Supplementary Online Material, SOM “Design of sensitivity benchmark”) measured the success in detecting homologs through a score referred to as *AUC1*, namely the fraction of annotated homologs found until the first non-homolog. Similar to other measures scoring search success, AUC1 depends heavily on the size of DB and the particular relation equated with homology, i.e., results differ between aiming at identifying pairs with similar structure, or similar function, or related in evolution, and given the immense diversity in definitions for function, AUC1-like measures can easily differ by an order of magnitude depending on the precise definition for function (Rost, 2002).

Alignments between pairs of sequences (also referred to as *pairwise* or *sequence-sequence*) unravel only simple connections, in evolutionary terms, the homology between proteins from closely related organisms. In order to intrude deeper into the *<u>twilight zone</u>* of sequence comparisons (Doolittle, 1986; Rost, 1999; Yona and Levitt, 2002), we need to find a family of related proteins through pairwise alignments, compile a profile or position-specific scoring matrix (PSSM) from this family, and then use the profile for more fine grained sequence-profile alignments (e.g. *Clustal* (Higgins and Sharp, 1989) or *PSI-BLAST* (Altschul et al., 1997)). The signal distinguishing between related and unrelated proteins becomes obfuscated upon entering the *twilight* zone; in fact, the transition from what we may call “*<u>daylight</u>*” to *twilight* zone is described by a rapid order-of-magnitude signal loss akin of a phase-transition (Rost, 1999). Profile-based searches intrude into the twilight zone, in particular, methods based on Hidden Markov Models (HMMs) as introduced for protein comparisons almost 30 years ago (Haussler et al., 1993; Krogh et al., 1994) and perfected through HHblits (Remmert et al., 2012) and Jackhmmr (Johnson et al., 2010).

Even more powerful than sequence-profile are profile-profile comparisons using profiles for query and database (Remmert et al., 2012). While some relations in this realm may no longer be indicative of evolutionary connections, at least many of the relations obtained by comparing the three-dimensional (3D) structures of proteins reveal that many relations are likely indicative of evolutionary connections so distant that even advanced sequence-based alignment methods fail to unravel those (Orengo et al., 2001; Nepomnyachiy et al., 2017). Thus, the identification of relations in the lower end of the twilight zone is often referred to as remote homology detection. Even profile-profile comparisons usually fail to intrude even further, namely the *<u>midnight</u> zone* in which sequences have diverged to random levels (5-10% pairwise sequence identity, <u>PIDE</u>) (Rost, 1997; Friedberg et al., 2000; Nepomnyachiy et al., 2017). In fact, most pairs of proteins with similar structures populate this realm (Rost, 1997; Friedberg et al., 2000).

Profile-based methods vary in speed, from the highly optimized *HHblits* (Remmert et al., 2012) averaging 2 min per query against UniRef30 (Remmert et al., 2012), to the lightning fast iterated *MMseqs2* profile-aligning in sub-seconds on UniRef90 (almost three orders faster than HHblits). Runtime details crucially depend on parameter choices such as “number of hits reported”.

### Protein Language Models (pLMs) capture crucial constraints

Faster computer hardware (largely GPUs and TPUs), better algorithms in Machine Learning, and big data combined to leap Natural Language Processing (NLP). In analogy, *<u>protein Language Models</u>* <u>(pLMs)</u> use large databases of raw protein sequences to implicitly understand the language of life (Alley et al., 2019; Bepler and Berger, 2019; Heinzinger et al., 2019; Elnaggar et al., 2021; Ofer et al., 2021; Rives et al., 2021). Indeed, many pLMs essentially needed access to sequence collections ten times larger than UniProt (Suzek et al., 2015; UniProt Consortium, 2017), namely BFD (Steinegger and Söding, 2018; Steinegger et al., 2019). Some pLMs additionally include supervised training (Bepler and Berger, 2019; El-Gebali et al., 2019). The values from the last hidden layers of the pLMs typically are extracted as “*<u>the embeddings of the pLM</u>*”. For the pLM ProtT5 (Elnaggar et al., 2021), in particular, these embeddings have 1024 dimensions for each residue in the protein (each protein position). The mean over all per-residue embeddings in a protein (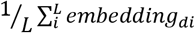, with L as the protein length and *embedding*_*di*_ as the embedding of residue *i* in dimension *d*) yields a new per-protein embedding (global average pooling) of the same dimension (1024d for ProtT5). We used this per-protein embedding as feature for our search.

Embeddings from pLMs capture information beyond sequence similarity and can help to detect close and remote homologs (Rao et al., 2019; Littmann et al., 2021a; Littmann et al., 2021b; Rives et al., 2021; Heinzinger et al., 2022). The similarity between protein-pairs in terms of embedding and sequence space are only weakly correlate which allows embedding-based annotation transfer (EAT) even for proteins with different sequences (PIDE<20%)(Littmann et al., 2020; Heinzinger et al., 2022). The per-residue embeddings as sole input to relatively shallow subsequent AI improve per-residue predictions of secondary structure (Elnaggar et al., 2021), inter-residue distance (Weissenow et al., 2022), 3D structure (Weissenow et al., 2022), and even residue-conservation and effects of sequence variation (Marquet et al., 2021; Dunham et al., 2022) beyond top prediction methods using evolutionary information from MSAs. Although falling substantially short of *AlphaFold2* (Jumper et al., 2021). Per-protein embeddings outperform the best MSA-based methods in the prediction of sub-cellular location (Staerk et al., 2021), signal peptides (Teufel et al., 2021) and binding residues (Littmann et al., 2021c).

### Nearest neighbor search through pLM embeddings

To search in embedding space, we want to find the k embeddings in a dataset most similar to our query given a distance metric. This is known as nearest neighbor search. As determining the exact nearest neighbors becomes intractable in high-dimensional spaces (Slaney and Casey, 2008), we applied approximate nearest neighbor search (k-nn) that is well established in domains including image recognition (Liu et al., 2007; Li et al., 2018), recommender systems (Bernhardsson, 2020) and NLP (Khandelwal et al., 2019). Modern indexing techniques such as Hierarchical Navigable Small World Graphs (Malkov and Yashunin, 2018) or Product Quantization (Jegou et al., 2010), as well as, approaches building upon those two (Babenko and Lempitsky, 2014; Baranchuk et al., 2018; Matsui et al., 2018) handle billion-scale datasets, suggesting the applicability to searching and/or clustering databases such as TrEMBL with 195M sequences (11/2020 used here) (UniProt Consortium, 2017) or even the entirety of BFD (Steinegger and Söding, 2018; Steinegger et al., 2019).

### Standard of truth: CATH and Pfam

We benchmarked on two databases: CATH (Orengo et al., 1997; Sillitoe et al., 2019) and Pfam (Bateman et al., 2000). CATH is created by three main steps: (1) parse all proteins of known 3D structure taken from the PDB (Protein Data Bank (Burley et al., 2017)) into compact domains. (2) Align all domains to each other by methods comparing 3D structures, i.e. structural alignment techniques (Orengo et al., 1992; Kolodny et al., 2005). (3) Proteins of unknown 3D structure are aligned by HMMer (Finn et al., 2011) to the 3D-aligned domain *seeds* forming the four classes of CATH: C (class), A (architecture), T (topology), H (homologous family). The Pfam database (El-Gebali et al., 2019) collects protein families without using 3D structure information. Consequently, Pfam family seeds are much shorter than structural domains (Liu and Rost, 2003), incidentally, also built using HMMer (Finn et al., 2011).

### Advantages of pLMs

The key advantage of pretrained pLMs is that might implicit capture the same constraints that shaped evolution. Could this same advantage be harnessed to also revolutionize sequence comparisons? Here we analyzed to which extent alignments using the generalized sequences as found in embeddings might be competitive with traditional sequence-based approaches.

## 2 Methods

### Data set 3D: CATH20

We used a redundancy-reduced version of CATH v4.2.0 (Orengo et al., 1997; Sillitoe et al., 2019) provided by the CATH team, which was optimized to contain as many sequences as possible. It was created by eliminating pairs with ≥20% PIDE and ≥60% overlap with the longest protein and consists of 14,433 domain sequences in 5125 families. We computed embeddings for both datasets with the python api of bio_embeddings v0.2.0 (Dallago et al., 2021). The full 14,433 domains served as target database (to search against with the query), and 10,874 domains from the subset of 1,566 families with more than one member served as queries (to search with). We deemed a result as correct if the top hit (excluding self-hits) belonged to the same Pfam/CATH family as the query.

### Data set 1D: Pfam20

In practice, many users will either not know or use single domains for their searches because as many as 80% of all proteins may have several domains (Liu et al., 2004), and because for the target database users would be limited to domain-based resources such as CATH (Orengo et al., 1997; Sillitoe et al., 2019) or Pfam (Bateman et al., 2000; El-Gebali et al., 2019). To flip this perspective: most expert users will likely use one of those two at some point and will have some idea about the composition of structural domains in their protein, in particular, given the AlphaFold2 predictions for hundreds of millions of proteins (Tunyasuvunakool et al., 2021) that simplify separating proteins into structural domains.

We proxied searches with full-length proteins through the Pfam-based dataset. To have enough proteins with matching domains without over-representing large families, we picked 20 domains from each Pfam family with at least 20 members and accumulated all of those proteins into a set dubbed Pfam20. For these, we retrieved the full-length proteins for each Pfam-region. This provided 313,518 proteins and the set of all Pfam domain annotations for each protein. The task was to find all proteins that have a Pfam domain from the same family as any of the Pfam-domains annotated for the full-length query. This sampling ensured each query to have at least 20 correct hits. We searched this set in all-against-all fashion, considering a query-hit pair as correct if the two had at least one Pfam annotation in common. For k-nn, we retrieved the 300 nearest neighbors for each query. As most queries had 20 correct hits, the AUC1 (area under curve until first incorrect match) fell between 0 and 20 of 20.

### Sequence alignment

MMseqs2 version 13 (Steinegger and Söding, 2017) served as state-of-the-art (SOTA) for combining speed and sensitivity. We searched with a sensitivity of 7.5 (-s 7.5) and accepted hits with E-values ≤ 10^4 and a prefilter limit of 300 hits. For the CATH20 set, these settings found the correct hit in all but 11 of the 14,433 queries.

### Protein Language Model (pLM) embeddings

SeqVec (Heinzinger et al., 2019) applies the bidirectional Long Short-Term Memory layer (LSTM) architecture of ELMo (Peters et al., 2018) architecture from NLP to proteins, yielding embeddings that are 1024 dimensional 32-bit float vectors. ProtTrans (Elnaggar et al., 2021) abbreviates a collection of pLMs all of which use a transformer architecture and have more free parameters than SeqVec (ProtBert 420M – with M for million, ProtAlbert 224M, ProtT5 3B – with B for billion, vs. SeqVec 93M) and were trained on the much larger BFD dataset (UniRef50 50M) (Steinegger and Söding, 2018; Steinegger et al., 2019; Elnaggar et al., 2020). While ProtBert has the same embedding dimensionality as SeqVec (d=1024), ProtAlbert has 4096-dimensional embeddings. From the ESM-series of pLMs (Rives et al., 2021), we benchmarked ESM and ESM1b with 670M and 650M parameters respectively and 1280 embedding dimensions, which were trained on the 250 million sequences. ProtT5 (Elnaggar et al., 2021), based on the Text-to-Text Transfer Transformer (T5, (Raffel et al., 2019)), is a model consisting of an encoder and a decoder, of which we used only the decoder. The original ProtT5 model was only pretrained on BFD (ProtT5 BFD), a later version was finetuned on UniRef 50 (ProtT5 XL U50). We used it in half precision mode (fp16) for speedier embedding generation without loss of accuracy (Elnaggar et al., 2021).

### Embedding-based clustering

#### Step 1: k-nn index

We constructed an HNSW index (Hierarchical Navigable Small World Graphs) of M=42 (Malkov and Yashunin, 2018), and searched with efSearch=256 using faiss (Johnson et al., 2019). While storing the embeddings for our datasets required 642 MB, the HNSW index, which included both the embeddings and the HNSW graphs, required 1.4 GB.

#### Step 2: k-nn score

As basis for the combined method, we used negative log-transformed E-values for the hits from MMseqs2 and cosine similarities for the embeddings. As we chose E<0.1 and since cosine scores were between 0 and 1, the transformed E-values were always larger than the cosine similarities, forming our new combined scored. However, for the raw E-values lower is better while for the k-nn scores the opposite held. The combined score used the same, more simplistic, normalization of “higher is better”.

### Hardware

All benchmarks were performed on a machine with 2 Intel Xeon Gold 6248 with a total of 80 threads, 400GB RAM and an Nvidia Quadro RTX 8000 with 46GB memory. The GPU was only used to generate embeddings.

### Implementation and availability

The k-nn search was performed using the python interface of faiss version 1.6.3 (Johnson et al., 2019). To align the k-nn hits, we wrote them into a MMseqs2 prefilter database and ran MMseqs2 align with an E-value cutoff of 10,000 (-e 10000). We made the code that reproduced all figures and tables as well as all figure data available at https://github.com/konstin/knn-for-homology, together with the raw data of the figures.

### Performance measures

#### Top1(CATH20)

For the CATH20 dataset, we searched each of the 10,874 domains from a family with more than one representative (query) against the 14,432 other domains in the CATH20 (database) with an exhaustive nearest neighbor search with cosine similarity. For each query, we considered the search result correct if the top hit was from the same homologous superfamily as the query. We considered only the top/first hit as it is has been suggested to be the most relevant for homology-based inference (Littmann et al., 2020). We reported two values reflecting performance accuracy. (1) Raw accuracy was defined as the fraction of correct hits:

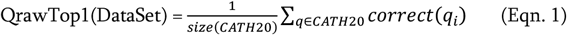

with DataSet as CATH20|Pfam20, size(DataSet) gave the number of queries in the data set, qi was the i-th query in DataSet. (2) As another measure, we considered the normalized accuracy normalized by family size to remove the bias from large families:

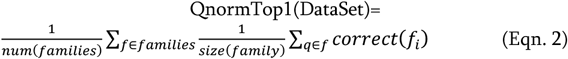

The latter was obtained by computing the accuracy for each family separately and taking the mean over these accuracies. These two scores have also been referred to as micro and macro average. For CATH20, we have reported both measures because the two often differed substantially. While the normalized accuracy removes the bias towards large queries, the raw accuracy adequately represents the abundance of sequences in the redundancy-reduced set.

#### AUC1(Pfam20)

For Pfam20 for each query, we recorded the fraction of true positive hits detected up to the first false positive. This sensitivity is also called area under the curve 1 (AUC1).

Errors were estimated using 500 rounds of bootstrapping and extracting the 95% confidence interval.

## 3 Results

### 3.1 Single domains: CATH (set *CATH20*)

#### Proof-of-principle: identification of domains

The embedding-based nearest neighbor search (k-nn) found more diverged/sequence-dissimilar homologous domains than *MMseqs2*, the state-of-the-art (SOTA) for fast and sensitive sequence-based search. Embeddings from more advanced pLMs clearly outperformed those from simpler pLMs (Fig. 1, Table 1). This finding is easiest to illustrate using “>” to mean “better than” for the major pLMs (Fig. 1), we observed: ProtT5>ESM1b>ProtAlbert> ProtXLNet>ProtBert. All differences were statistically significant at the 95% confidence interval (CI). While *MMseqs2* outperformed the less advanced pLMs, it was outperformed by ProtT5 and ESM1b (Table 1). SeqVec’s LSTM1 layer performed better than any combination of three layers (Supporting Online Material (SOM), Fig. S3).

**Fig. 1:**
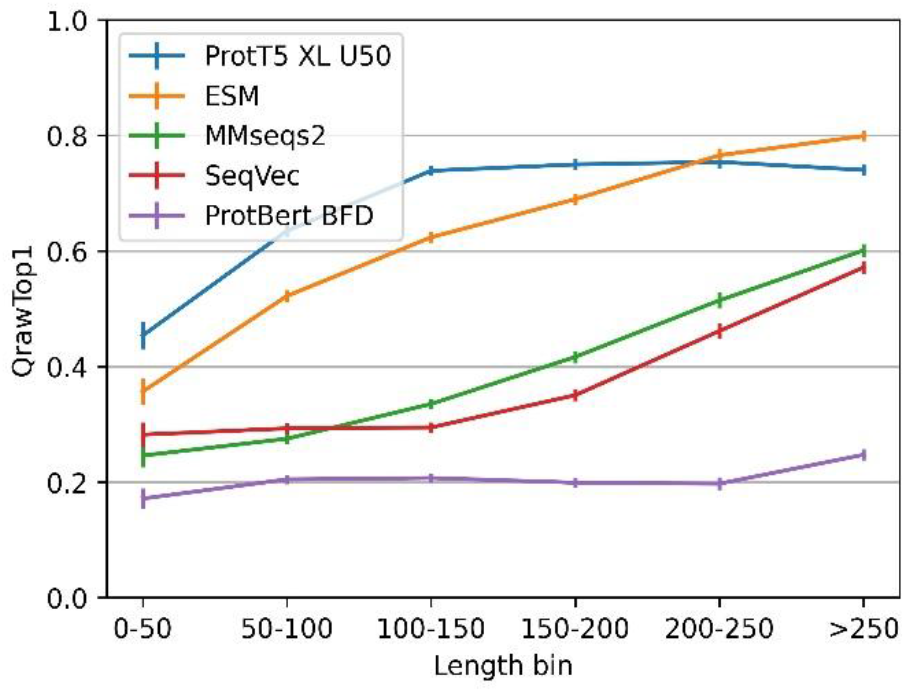
Performance better for longer proteins. The vertical y-axis QrawTop1 (Eqn. 1) reflect the performance for proteins from the length interval specified on the horizontal x-axis (non-cumulative bins). While the embeddings from ProtT5 and ESM saturated quickly, *MMseqs2* and the *SeqVec*-embeddings correlated more linearly with protein length.

**Table 1:**
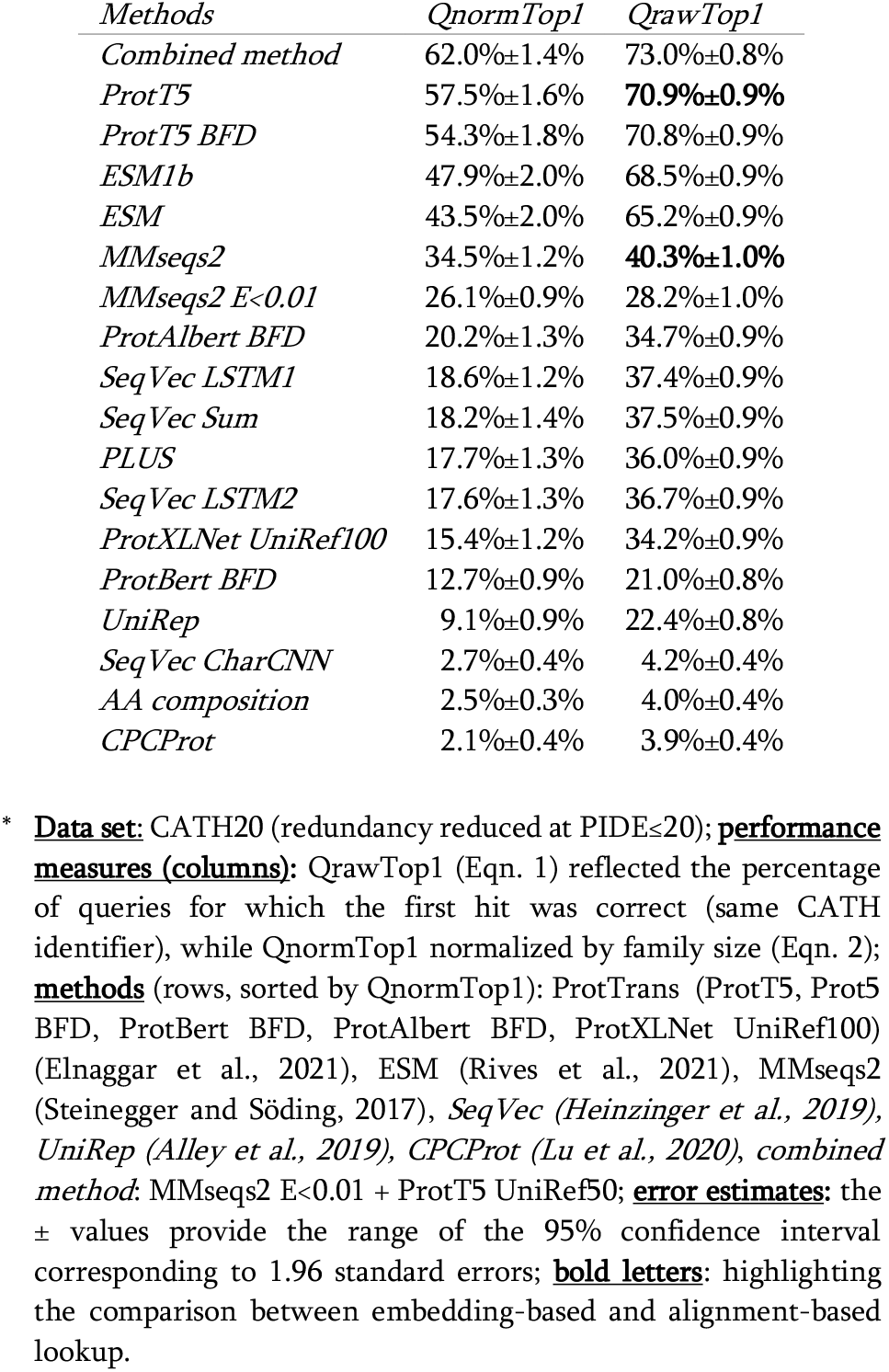
Performance on CATH20. *.

In terms of embedding measure (proximity/distance of two embeddings vectors of, e.g., 1024 dimensions for ProtT5, representing query and database hit), the cosine similarity consistently outperformed the Euclidean distance, albeit only slightly (Table S1). For instance, the normalized accuracy (QnormTop1) for ProtT5 dropped from 57.5%±1.6% (using cosine similarity to measure that the query-hit vector are similar) to 55.3%±1.7% (using the Euclidean distance to measure the embedding similarity).

To clarify the novel contribution of embeddings, we replaced all hits from *MMseqs2* with E-value>T and those with no match (11 cases) with hits from *knnProtT5*. The resulting combined search results (dubbed “*combined method*”) outperformed both methods over a wide range of thresholds T (Fig. S1, Fig. S2). For instance, at E-values of T=0.01, the raw accuracy increased by over two percentage points to 73.0% (Table 1, first row). The combined method outperforms both methods’ accuracy over a large range of cutoffs (Fig. S1 and Fig. S2).

#### Hypothetical best of both

If we could by some unknown procedure pick only the correct hits from each method, we would reach QrawTop1=78.2%. This implied a higher increase from *combined method* to *hypothetical* than from *knnProtT5* to *combined method*, but much less than from *MMseqs2* to *combined*. This hypothetical perfect merger marked the theoretical limit for the combined approach, which implied that the simple E-value threshold of T=0.01 already reached almost half of the improvement theoretically possible.

#### Factors determining performance

To better understand the complementarity of alignment-based and embedding-based approaches, we zoomed into strengths and weaknesses of each approach. For *MMseqs2* and *SeqVec* accuracy clearly correlated with protein length (“shorter proteins better”), while for ProtT5 and, to a slightly lesser extent, for ESM, the accuracy saturated for longer proteins. ProtBert-BFD, on the other hand, performed rather consistently across the spectrum of protein length (Fig. 1). By design, CATH20 considered only single domains. We observed only a limited correlation between cosine similarities and E-values with a Spearman’s ρ of −0.17 and Pearson Correlation Coefficient between cosine similarities and the log-transformed E-values of −0.14. This confirmed prior results (Littmann et al., 2020).

### 3.2 Full-length proteins: Pfam regions (set *Pfam20*)

#### Embedding-based *knnProtT5* alone not competitive

The above assessment focused on the comparison of single-domain proteins or single domains. The *Pfam20* (Methods) benchmark pulled in full-length proteins (still compared to single domain-like Pfam regions). Instead of outperforming *MMseqs2* (for *CATH20*), *knnProtT5* performed clearly worse for *Pfam20* (Fig. 2: solid red and dotted green below solid blue; AUC1(knn,Pfam20)=0.367 vs. AUC1 (Mmseqs2,Pfam20)=0.52). While *knnProtT5* and *MMSeqs2* found a similar fraction of homologs in the first 300 hits (68.0% vs. 67.4%), the vector distances (cosine similarity) between the per-protein representations did not sort the hits precisely enough, leading to a drops in AUC1 specifically and on the entire precision-recall more generally (Fig. 3; Fig. S5).

**Fig. 2:**
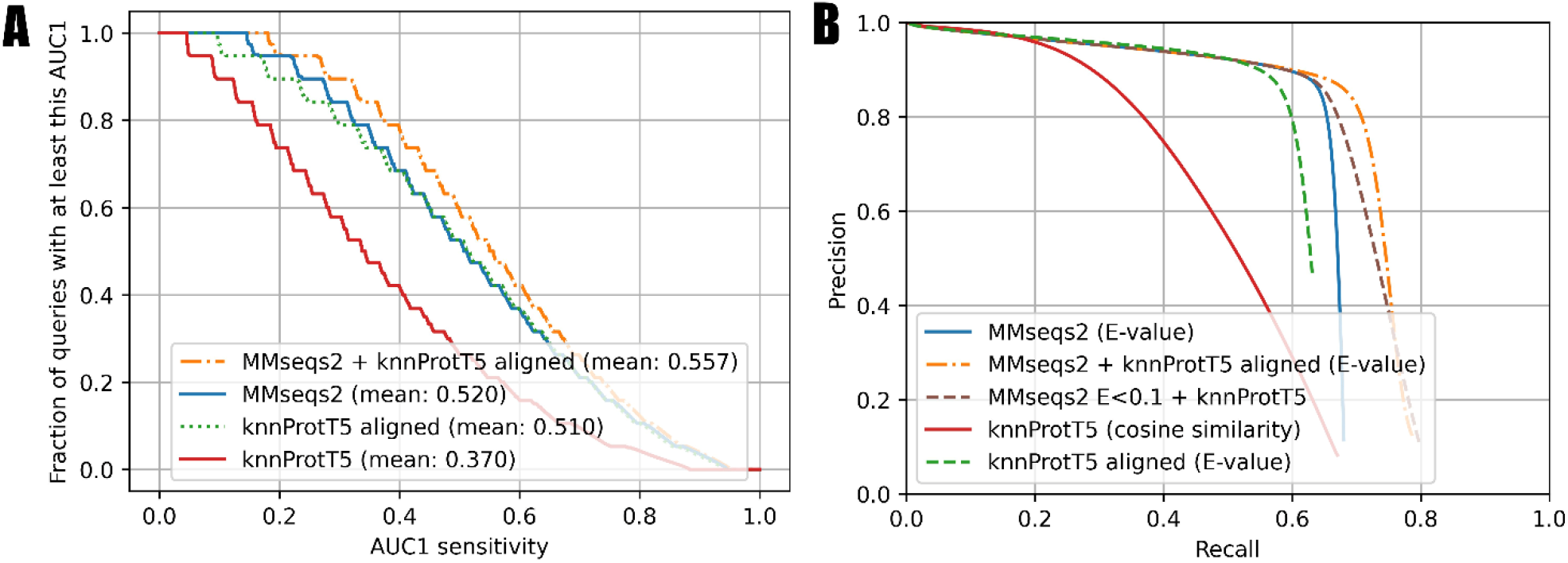
Sequence-based approaches better for full-length proteins (Pfam20). <u>Data set</u>: *Pfam20;* we searched all-against-all through 313,518 proteins with Pfam (Bateman et al., 2000) annotations. <u>Panel A:</u> Cumulative distribution of AUC1 that for each query reflected the fraction of queries (against the 313,518) with a matching Pfam annotation ranked above the first hit without matching Pfam annotation. Higher curves implied higher sensitivity. The steps in the lines originated from sampling 20 members for each Pfam family. *MMseqs2* (Steinegger and Söding, 2017) as state-of-the-art in fast and sensitive sequence-based alignment in solid blue; the novel *knnProtT5* method in solid red; *knnProtT5* for search plus Smith-Waterman alignment performed by *mmseqs align* in dotted green; the combination of *MMseqs2* and *knnProtT5* in dot-dashed orange. <u>Panel B:</u> Precision-recall curve for the same methods as in panel A except for the dashed green marking an additional test combining unaligned *knnProtT5* hits with *MMseqs2* to show the power of embeddings on their own. The two combined approaches use MMSeqs2 E-values as base and *knnProtT5* with cosine similarity only where *MMseqs2* had low E-values. This explains the large overlap with *MMseqs2*.

**Fig. 3:**
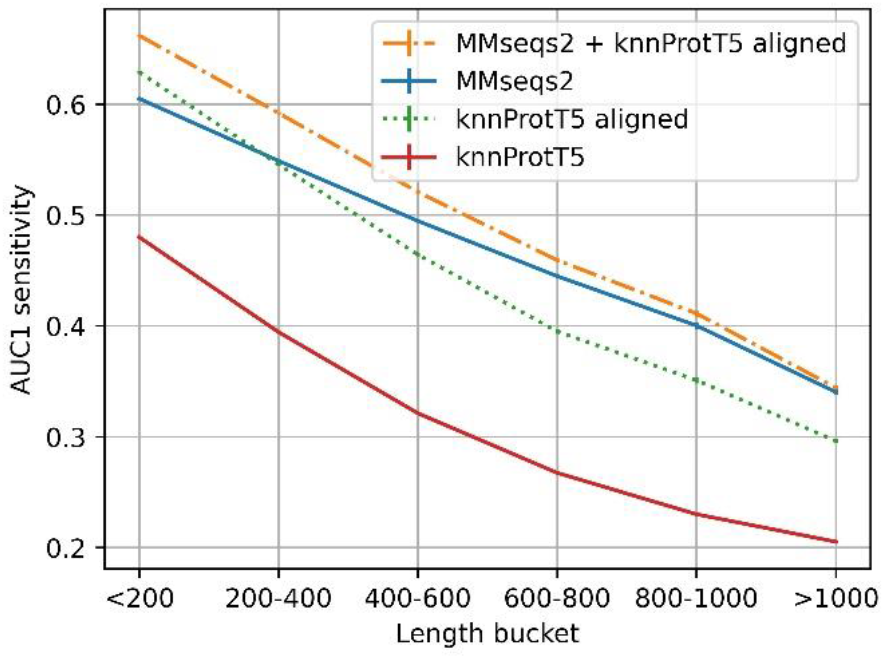
Longer proteins more difficult for Pfam20. Mean AUC1 sensitivity for different bins of lengths (number of residues) for the query protein (full proteins, not just domains compared to Pfam regions), showing how the combined method works across different sequence length. For those full-length proteins, all methods performed better for shorter than for longer proteins, e.g., MMseqs2 performance was almost half for proteins shorter than 200 residues than for those longer than 1000 residues, and still substantially better for proteins with <600 residues that account for the majority of UniProt. Long proteins are often contain multiple domain which have more total homologs on average and are also more difficult to match on per-protein representation level (Discussion).

#### Shuffling domains

By using Pfam annotations as ground truth, we might incorrectly consider a hit as incorrect when domain annotations are missing. The common solution is to shuffle the amino acid sequence outside of domain annotations and/or to add reversed or shuffled sequences as known incorrect hits (Brenner et al., 1998; Steinegger and Söding, 2017; Buchfink et al., 2021). However, we found that ProtT5 clearly separated between reversed or shuffled and real sequence (Fig. S7, Table S2). Thus, we had to accept that some correct hits may be labeled as incorrect.

#### Combined method most sensitive

As for single-domains, the *combined method* (MMseqs2 + knnProtT5) considerably increased the overall sensitivity (Fig. 2B: *combined method*). Toward this end, we aligned the top 300 *knnProtT5* hits using MMseqs2 and merged those 300 with the top hits from *MMseqs2* ranked by E-value. This combination raised the AUC1 from 0.52 to 0.557 (Fig. 3) and improved the recall even for high precision (Fig. 3). The number of hits was the main hyperparameter for *knnProtT5* and the subsequent alignment. We chose 300 to be identical to the MMseqs2 prefilter default. Although for the embeddings of some pLMs the value mattered more, essentially all embeddings appeared relatively stable for choices above 150 hits (Fig. S6). In particular, the *combined method* clearly outperformed *MMseqs2* choosing the top 600 rather than 300 hits (AUC1(300)=0.520 vs. AUC1(600)=0.523). Thus, more hits correlated with increased AUC1 at the expense of runtime (below).

We also combined *MMseqs2* and *knnProtT5* without aligning the k-nn hits. Toward this end, we first filled the result list for a query with the *MMseqs2* hits at E-values<0.1 and then appended the k-nn hits, i.e., we accepted very reliable *MMseqs2* hits, and added *knnProtT5* hits to fill up the hitlist to 300. This simpler scheme also reached an AUC1 of 0.558, but the recall only improved for lower precision (Fig. 2B). The results were very similar to those for the *combined method* with re-alignment, reaching the same AUC1 of 59%.

#### Aligning *knnProtT5* hits competitive

Aligning the *knnProtT5* hits with Smith-Waterman (Smith et al., 1981), yielded an AUC1 similar to that of *MMseqs2*. Using *knnProtT5* as prefilter instead of *MMseqs2* is, however, infeasible in practice due to the immense amount of time needed to compute per-protein embeddings. Although we aligned with an E-value cutoff of 10000, the mean recall over all up to 300 hits dropped from 67.4% to 63.8%, or put differently: 3.6% of all homologs were correctly found by *knnProtT5* but then dropped because they could not be aligned adequately.

#### Lower AUC1 for longer proteins

While *knnProtT5* correctly retrieved many hits for long proteins, barely any of those were not already found by MMseqs2 (Fig. 3). Splitting long proteins into overlapping slices of 600 residues, which were embedded individually and searched all-against-all against a databases of slices, did not improve (results not shown). Due to the quadratic growth in terms of costs (time and memory) for computing ProtT5 embeddings for longer proteins, we had to remove 0.6% of the proteins with over than 3096 residues, worsening the results for the >1024 bucket slightly (Fig. 3). We observed that long sequences had disproportionally many hits with high cosine similarity (>0.95) and no matching annotations, most likely due to missing annotations.

#### Runtime: method fast, but slowed down by embedding-lookup

MMseqs2 took 17min 39s for the search (12m 2s prefilter and 5m 37s align). Generating embeddings took 7h 23min, giving an average of 0.08s per protein. Generating a Hierarchical Navigable Small World Graph (HNSW (Malkov and Yashunin, 2018) took 15s, the search took 77s. Compared to exhaustive nearest neighbor search we lost 0.004 AUC1 sensitivity (0.367 to 0.371), while the effect was below standard error for aligned *knnProtT5* and the *combined method*.

## 4 Discussion

### Key step: comparing generalized sequences

Embeddings from protein Language Models (pLMs) appear to carry information about aspects such as protein function, structure, and evolution (Rao et al., 2019; Littmann et al., 2020; Elnaggar et al., 2021; Littmann et al., 2021a; Littmann et al., 2021b; Marquet et al., 2021; Rives et al., 2021; Dunham et al., 2022; Heinzinger et al., 2022; Weissenow et al., 2022). In this sense, they constitute what we might refer to as “generalized sequences”. The key advance underlying our novel approach is to use generalized sequences for remote homology detection. While the idea for this is not new (Alley et al., 2019; Littmann et al., 2021a; Littmann et al., 2021b; Ofer et al., 2021; Bileschi et al., 2022; Heinzinger et al., 2022; Nallapareddy et al., 2022), here we presented a more rigorous and generic framework for directly comparing embedding-based to sequence-based alignments which have been optimized which have been optimized for half a century. Despite this advantage of being decades ahead in experience, methods that train on embeddings to map proteins to particular databases, such as CATH (Orengo et al., 1997; Sillitoe et al., 2019) and Pfam (Bateman et al., 2000) already outperform traditional sequence-based methods, even those based on profiles (Bileschi et al., 2022; Heinzinger et al., 2022; Nallapareddy et al., 2022). Here, we explored to which extent pLM embeddings directly, i.e., without any further training, are competitive in terms of performance and speed to a state-of-the-art (SOTA) sequence-based identification of homologs.

Our approach of finding k-nn embedding matches appeared to have several advantages. Firstly, by using k-nn matches (dubbed *knnProtT5*), we could explicitly drop the incorrect and limiting assumption that alignments at position P1 are statistically independent of those at position P2. Secondly, by replacing an amino acid alphabet with a vector condensing information from other residues, possibly far away in terms of sequence separation, that influence the evolution, function and structure at each residue position P, we implicitly use such constraints to compare sequences. Thus, although other novel solutions for non-iterated homology searches based on embeddings tend to focus on speed, our solution tried to combine speed and sensitivity. This was most prominently exemplified by the 3.6% of ground truth homologs which *knnProtT5* found but which could not be reasonably aligned (neither by *MMseqs2* (Steinegger and Söding, 2017) nor by Smith-Waterman (Smith et al., 1981)).

Overall, raw embeddings through k-nn (*knnProtT5*) could improve over traditional sequence similarity searched, both for single-domain vs. single-domain homology-based inference (CATH20, Fig. 1) and for more general full-length vs. single-domain/Pfam-region homology searches (Pfam20, Fig. 2). Merging sequence- and embedding-based (MMseqs2 + *knnProtT5*) did better than any of the two throughout (Fig. 1-2: *combined method*). However, for the more realistic use-case of comparing full-length proteins against Pfam-regions, MMseqs2 overall clearly outperformed the raw embeddings (Fig. 2). Thus, we established an idea for a simple solution of using embeddings without further machine learning that showed some promises and strengths without breaking through.

### Speed not necessarily sufficient

While the nearest neighbor search itself is blazingly fast, the time needed to generate the per-protein embeddings by ProtT5 and the effective GPU requirement might throw up major hurdles for adopting our approach. In fact for full-length proteins, the iterated profile search from MMseqs2 currently yields better results in shorter time. Two future eventualities might change this: firstly, databases such as UniProt (UniProt Consortium, 2017) might offer per-protein embeddings for all their proteins. If so, *knnProtT5* would immediately become competitive. Secondly, judging from advances in NLP, we expect significant model speedups, potentially making k-nn with alignment viable on its own. PLMs leaped through rounds of exponential improvements over the last three years: from embedding-based prediction methods being faster than multiple sequence alignment (MSA) based predictions but much worse to outperforming MSA-based methods. Given this rapid evolution, large pLMs may soon become sufficiently good and/or small (NLP on smart phone) to justify the runtime cost. For instance, only the more recent ESM (Rives et al., 2021) and ProtT5 (Elnaggar et al., 2021) outperformed *MMseqs2* for CATH20, while slightly earlier models such as SeqVec, or ProtBERT failed to do so. Maybe the next leap for the next pLM by increasing power and reducing the size will shift the balance more toward embedding- than sequence-based solutions.

### Multiple domains will continue to challenge comparisons of entire proteins

As many as 80% of all proteins may consist of several domains. If we chopped all proteins into their domains and compared all-domains-against-all, the then full-domain embedding based k-nn succeeded (Fig. 1), while for the comparison of full-length proteins against domains, the average-pooled per-protein embeddings of the queries are too coarse-grained (Fig. 2). Sequence-based solutions built upon the local Smith-Waterman concept (Smith et al., 1981) still succeed because of matching subsequences. In fact, this most likely explained the lack of improvement for proteins longer than 1,000 residues (Fig. 3). Another possible path might be to directly create pLMs capturing entire domains as the “units”, however, so far there seems no solution in this direction that succeeded without having to retrain specific AI models on top of embeddings, such as CATHe (Vaswani et al., 2017) or ProtTucker (Heinzinger et al., 2022), or the Pfam-AI (Bileschi et al., 2022).

While only tangentially relevant for homology search, we considered ProtT5’s ability to detect “fake” sequences and make them trivially separable a noticeable result in its own right.

### No advance without alignment?

A fundamental strength and limitation of our approach is that k-nn hits need to be aligned, e.g., by Smith-Waterman (Smith et al., 1981). Alignments become unreliable at low levels of sequence identity, i.e., exactly in the realm for which embedding similarity promises to be useful (Rost, 1999; Steinegger and Söding, 2017; Littmann et al., 2020; Buchfink et al., 2021). Indeed, *knnProtT5* found many correct hits below 20% PIDE that could not be aligned correctly (Fig. S6: *knnProtT5* vs. *knnProtT5*_*aligned*). These results might suggest the need for the development of an embedding-based local alignment method to use the full potential of embedding based homology search and to make hits interpretable beyond a single score. A few approaches have been proposed toward this end, however, these are limited to global alignments (Bepler and Berger, 2019; Morton et al., 2020), i.e., will even worsen the decline from domain vs. domain (CATH20, Fig. 1) to full-length vs. domain (Pfam20, Fig. 2).

### k-nn index for fast pre-filtering?

An exhaustive k-nn index search has quadratic complexity, making it unworkable for large datasets, compared to modern indices with log-linear runtime complexity (𝒪(n·logn)). The HNSW index we chose operated considerably faster than the *MMseqs2* prefilter while finding a comparable number of correct hits. It scaled well to billions of vectors (Malkov and Yashunin, 2018), making it feasible to search even metagenomics databases such as BFD (Steinegger and Söding, 2018; Steinegger et al., 2019). Combined with a fast pLM, this has the potential to outperform and outscale k-mer based approaches.

## 5 Conclusions

We demonstrated nearest neighbor search with protein Language Model (pLM) embeddings (here *knnProtT5*) to constitute a valuable complement to *MMseqs2*, enabling more sensitive homology searches. The embedding-based solution exceled in detecting distantly related domains (*remote homology detection*), finding hits that previously were not accessible by non-iterated homology search. Thus, embedding-based solutions might offer a more stringent baseline for the reach of homology based inference (if SIM(Q,A) > threshold, copy annotation from A to Q). *knnProtT5* also scales to increasingly large databases. The only limitation at the moment, the generation of per-protein embeddings, might be removed as an obstacle through database resources such as UniProt providing such embeddings in prestored fashion for all proteins. Either way, rapid progress of pLMs already renders nearest neighbor searches on embeddings a promising new path, allowing us to tap into a new pool of homologs from embedding space and to go beyond sequence similarity.

## 7 Author Contributions

KS, MH, MS and BR conceived and planned the experiments. KS carried out the experiments. KS wrote the manuscript with MH and BR. MS contributes MMseqs2 benchmarking.

## 9 Acknowledgments

Thanks to Tim Karl (TUM) for help with hardware and software; to Inga Weise (TUM) for support with many other aspects of this work. Christian Dallago (TUM) leading bio_embeddings and feedback throughout this work. Last, not least, thanks to all those who deposit their experimental data in public databases, and to those who maintain these databases.

## 10 Data Availability Statement

The datasets analyzed for this study can be found in the Pfam database (http://pfam.xfam.org/) and CATH database (https://www.cathdb.info/). Embeddings and intermediate databases are available through Zenodo under the doi 10.5281/zenodo.6794234.

## Abbreviations used

3D: three-dimensional
AI: Artificial Intelligence
BFD: Big Fantastic Database (Steinegger and Söding, 2018; Steinegger et al., 2019)
CATH: hierarchical classification of protein 3D structures in Class, Architecture, Topology and Homologous superfamily (Orengo et al., 1997; Sillitoe et al., 2019)
DL: Deep Learning
EAT: Embedding-based Annotation Transfer
EI: evolutionary information
embeddings: fixed-size vectors derived from pre-trained pLMs
ESM-1b: pLM from Facebook dubbed Evolutionary Scale Modeling (Rives et al., 2021)
FNN: Feed-forward Neural Network
HMM: Hidden Markov Model
HMMer: particular method for HMM-profile alignments (Finn et al., 2011)
LM: Language Model
MMseqs2: fast database search and multiple sequence alignment method (Steinegger and Söding, 2017)
MSA: Multiple Sequence Alignment
NLP: Natural Language Processing
PDB: Protein Data Bank (Burley et al., 2017)
Pfam: Protein family database (Sonnhammer et al., 1997; Bateman et al., 2000)(El-Gebali et al., 2019)
PIDE: percentage pairwise sequence identity
pLM: protein Language Model
ProtBERT: pLM (Elnaggar et al., 2021) based on the LM BERT
ProtT5: pLM (Elnaggar et al., 2021) based on the LM T5
PSSM: position-specific scoring matrix (also dubbed profile)
SOTA: state-of-the-art

