## Supplementary Online Material for "Nearest neighbor search on embeddings rapidly identifies distant protein relations"

### Supporting online material for: Nearest neighbor search on embeddings rapidly identifies distant protein relations

### Table of Contents for Supporting Online Material

### Short description of Supporting Online Material

In the main, we show the combined MMseqs2 + ProtT5 k-nn method on CATH20 only for an example cutoff of E<0.1 (Steinegger and Söding, 2017, Elnaggar et al., 2021, Orengo et al., 1997). Here we add figures for the whole range of E-value (Fig. S1 and Fig. S2 for *QrawTop1* and *QnormTop1*). Fig. S3 show how unlike the more complex linear combinations in ELMo (Peters et al., 2018), SeqVec for us performs best best when just using the hidden state from the LSTM1 layer. Table S1 shows how the cosine similarity beats the euclidean distance consistently. In Fig. S4, we confirm the low correlation between E-Value and cosine similarity.

For the Pfam20 dataset (El-Gebali et al., 2019), we show the correlation between cosine similarity and accuracy (Fig. S5) and how the number of hits we allow in a prefilter influences the fraction of homologs retrieved (Fig. S6). We can't shuffle unannotated regions in proteins as other benchmarks did (Steinegger and Söding, 2017, Buchfink et al., 2021) since ProtT5 can trivially separate real and fake sequences (Following section, Fig. S6 and Table S2)

---

##### T5 Detecting Shuffled Sequences

Real sequences vs. shuffled sequences ProtT5 can tell real sequences from reversed and shuffled sequences, generating considerably different embeddings. This can be shown by even only looking at principal component 1 of a PCA of the embeddings of native proteins and scrambled counterparts. As an example, we took 10000 random proteins from the Pfam set, generated embeddings for each original sequence, each sequence randomly shuffled and each sequence reversed. We project original and shuffled embeddings as well as original and reversed embeddings in a PCA. Supplementary Fig. S7 shows the distribution on embeddings across Principal Component 1. Using a cutoff of 0 with the values from this axis, this can be seen as a basic predictor for real vs. scrambled, for which we show confusion matrices in Supplementary Table 2. Surprisingly, we even get a good split without perform any kind of transformation by only taking embedding dimension 519, marking everything below 0.15 as real and everything above as scrambled. This alone reaches an accuracy of over 85% for each class on the tested set. This shows that ProtT5 can differentiate between real and "fake" sequences and that "fakeness" is the primary signal. This is also not due ProtT5 misses the methionine at the beginning of a protein as the same behaviour occurs with the domains from the CATH dataset.

### Material

**Fig. S1: CATH20 raw accuracy by E-value cutoff**

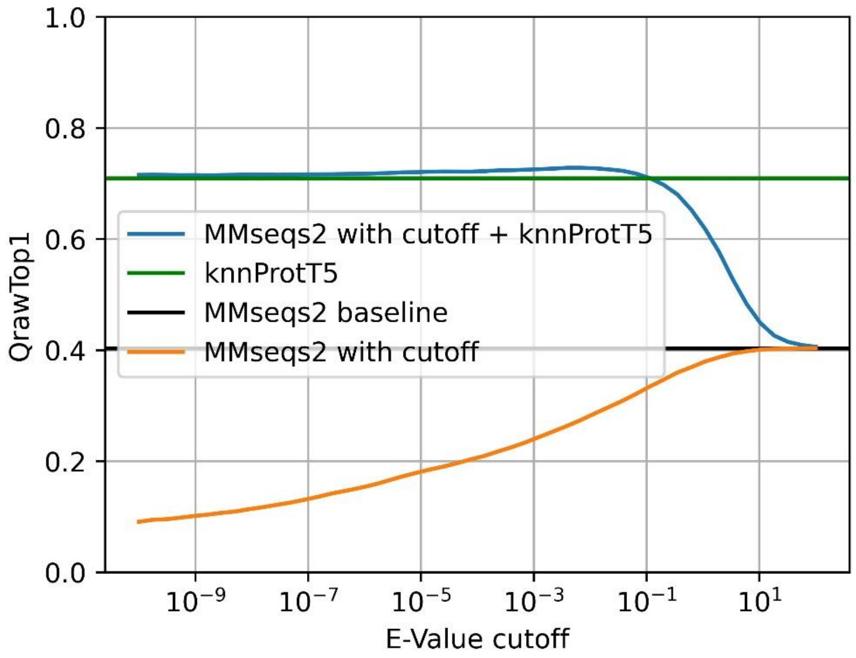

**Fig. S1: CATH20 raw accuracy by E-value cutoff.** The QrawTop1 accuracy of MMseqs2 {Elnaggar, 2021 #66} and the combined approach at different E-value cutoffs when searching for domains from the same homologous superfamily in CATH {Orengo, 1997 #74;Sillitoe, 2019 #49}. The blue line is the novel combined method, the green line is the proposed nearest neighbor search with ProtT5 XL U50 {Elnaggar, 2021 #66} and the black line is the MMSeqs2 baseline. The orange line is what we would get if we took the combined method and removed all the k-nn hits, which shows which fraction of the combined method score comes from MMseqs2 and which from k-nn. Equivalent of Fig. 2 with raw accuracies.

**Fig. S2: CATH20 normalized accuracy by E-value cutoff**

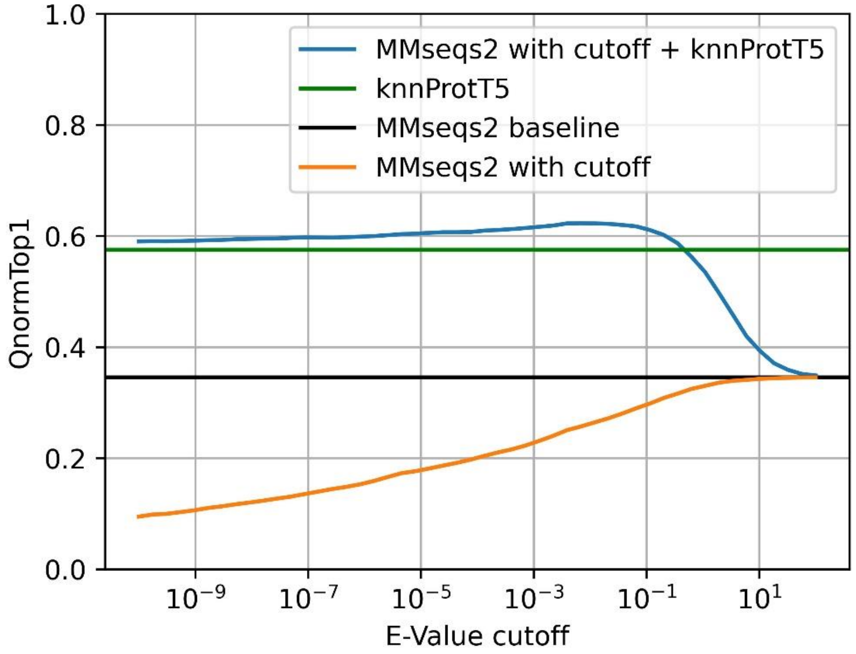

**Fig. S2: CATH20 normalized accuracy by E-value cutoff.** Identical to Fig. S1, except with QnormTop1 instead of QrawTop1

**Fig. S3: SeqVec layers for CATH20**

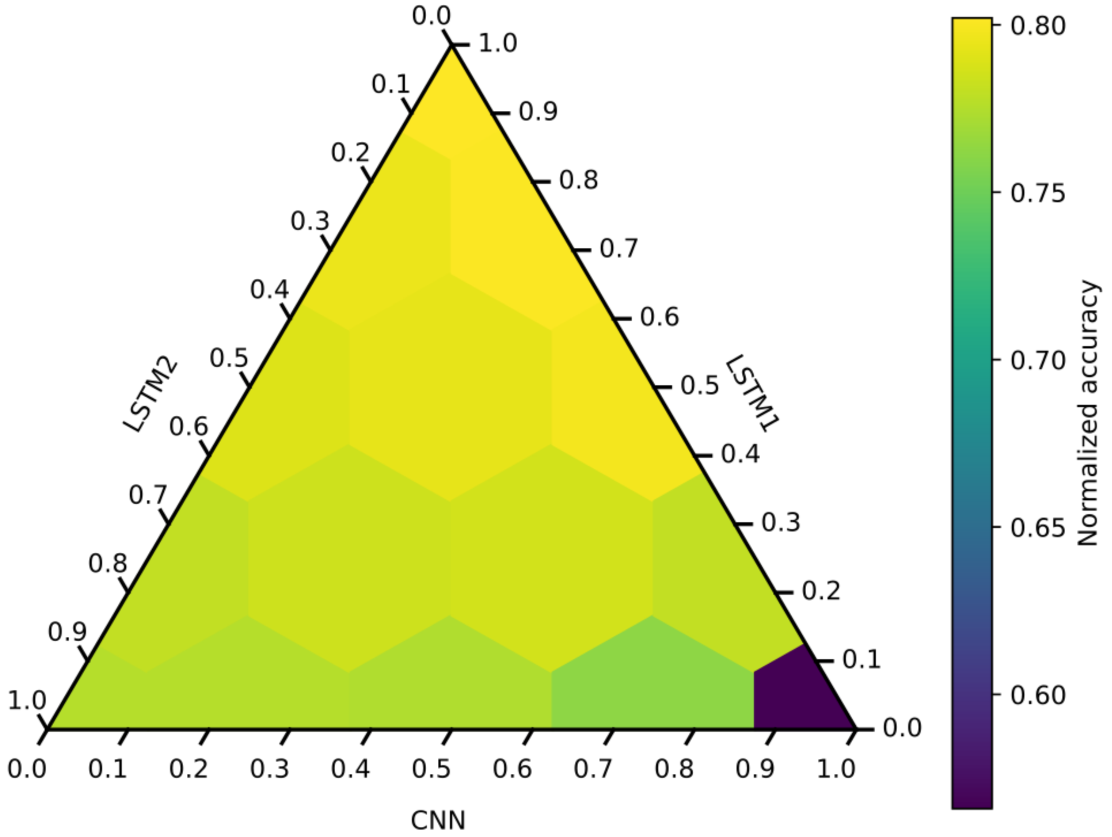

**Fig. S3: SeqVec layers for CATH20.** QnormTop1 on the CATH dataset for different linear combinations of SeqVec layers as ternary plot, showing how LSTM1 alone is the best layer choice, while combinations of layers do not improve over it.

**Fig. S4: CATH20 E-Value vs. Cosine Similarity**

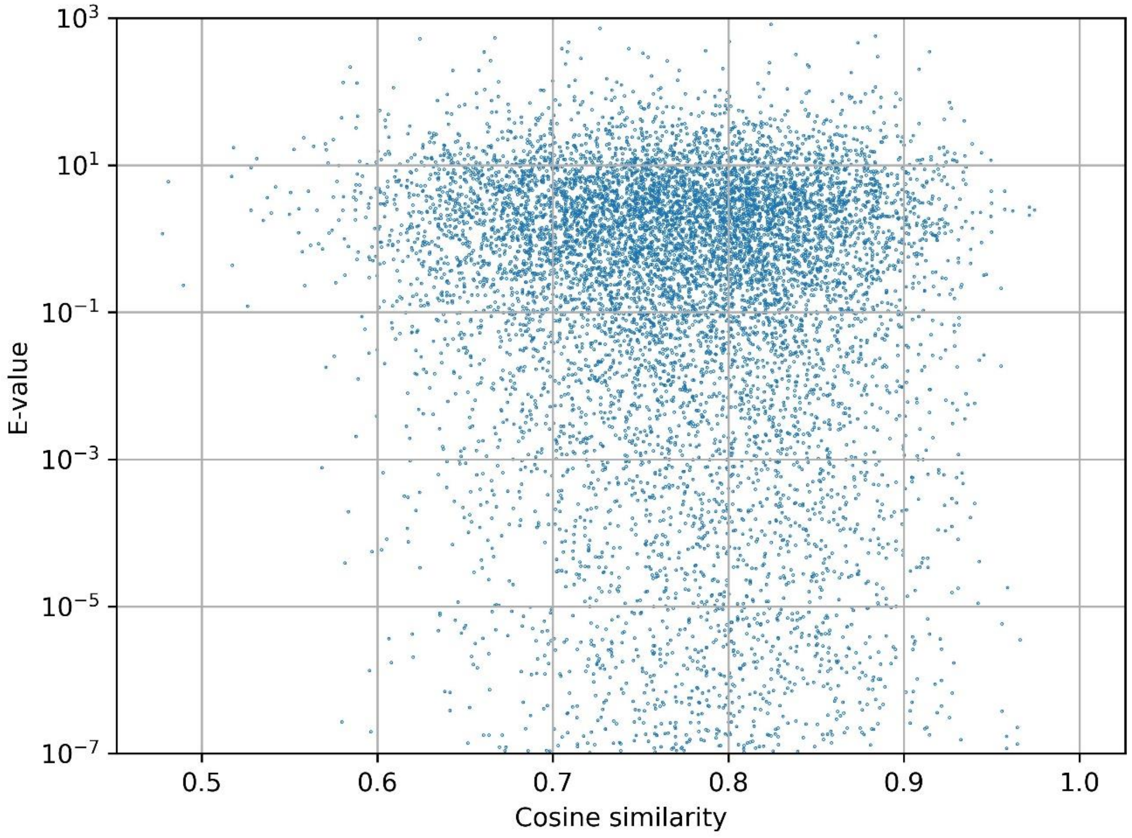

**Fig. S4: CATH20 E-Value vs. Cosine Similarity.** For each CATH20 query, we compare E-Value (MMseqs2) and cosine similarity (knnProtT5), showing the low correlation between the two scores

**Fig. S5: Pfam20 accuracy by cosine similarity cutoff**

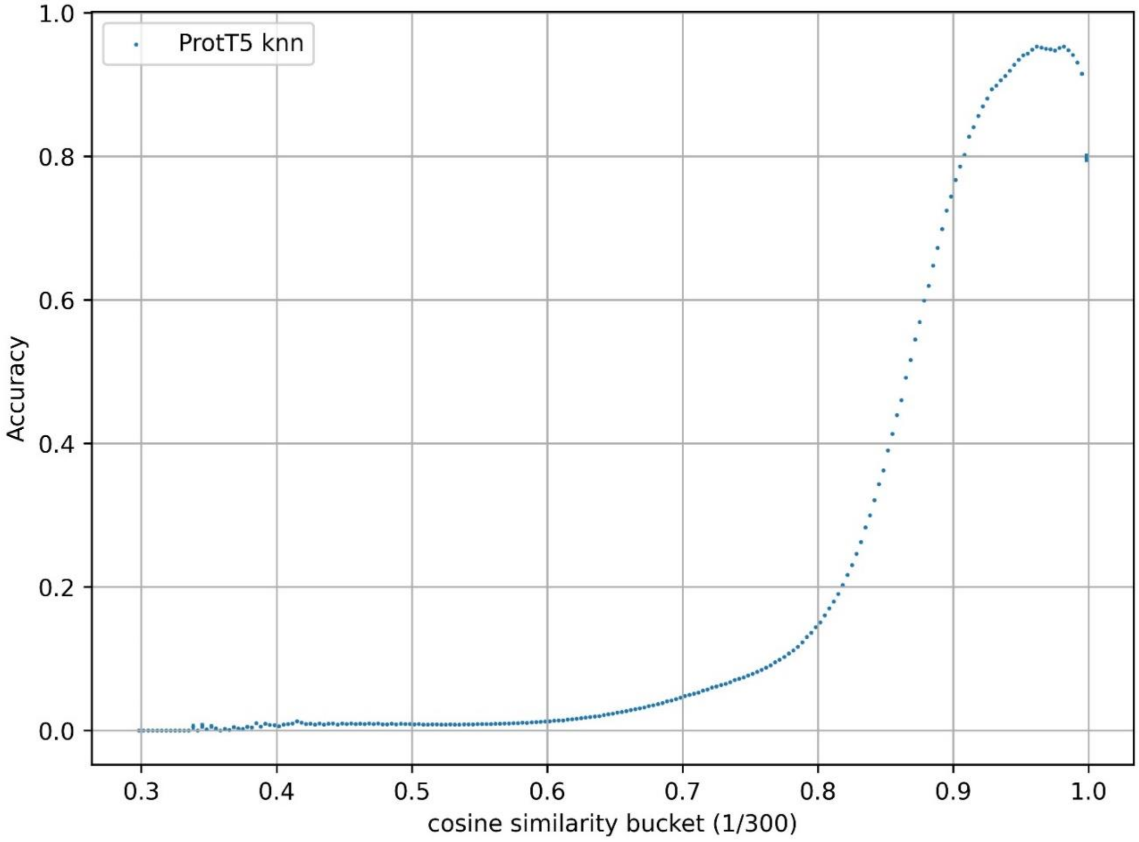

**Fig. S5: Pfam20 accuracy by score cutoff**. Accuracy of the knnProtT5 hits by their cosine similarity in the Pfam20 benchmark, displayed in bins of 0.01. The counterintuitive decline near 1 is due to missing annotations in Pfam: Of the 5233 matches with cosine similarity >0.95 marked as incorrect, 4577 were also found by MMseqs2 at E<0.0001, indicating they are most likely actually homologs not tracked in Pfam-A. We cannot avoid those unannotated yet correct matches since the common solution of shuffling/reversing unannotated regions is incompatible with language models (Fig. S7).

**Fig. S6: Influence of number of hits for Pfam20**

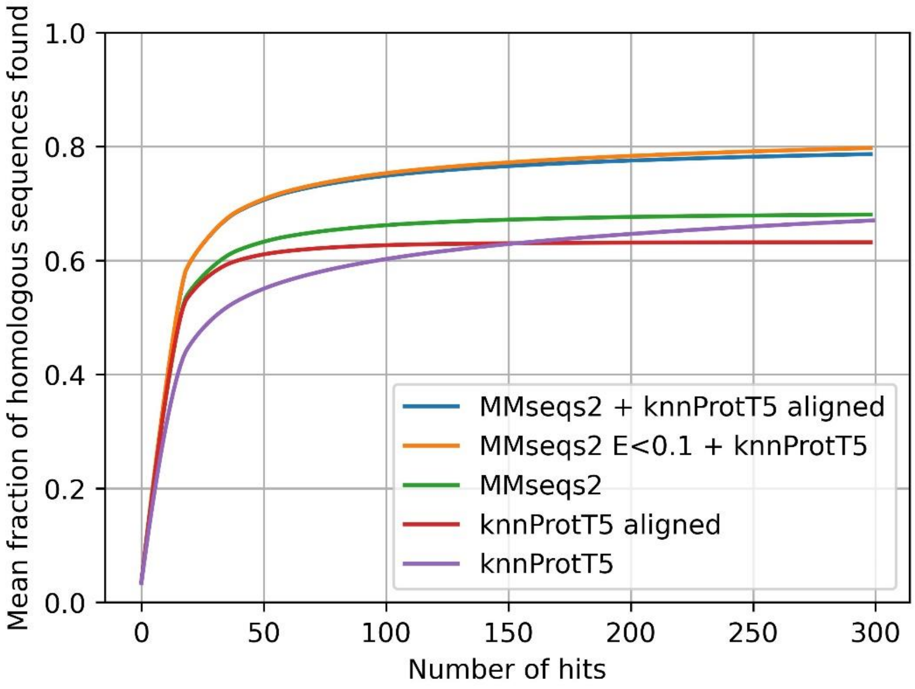

**Fig. S6: Influence of number of hits for Pfam20** The fraction of homologs recovered in the Pfam benchmark when considering a certain number of hits. ProtT5 k-nn alone is bad at sorting good hits to the front, even though it recovers about the same fraction of homologs as MMseqs2 after 300 hits. After alignment, most homologs are in the first few hits, however we use some homologs in the alignment process. Simultaneously, the number of hits is the most important hyperparameter in k-nn and for a following alignment; Picking e.g. only 50 hits from ProtT5 k-nn before alignment/combining would considerably impair performance. It also shows how the combined methods outperform the others consistently.

**Fig. S7: Comparing shuffled and reversed sequences**

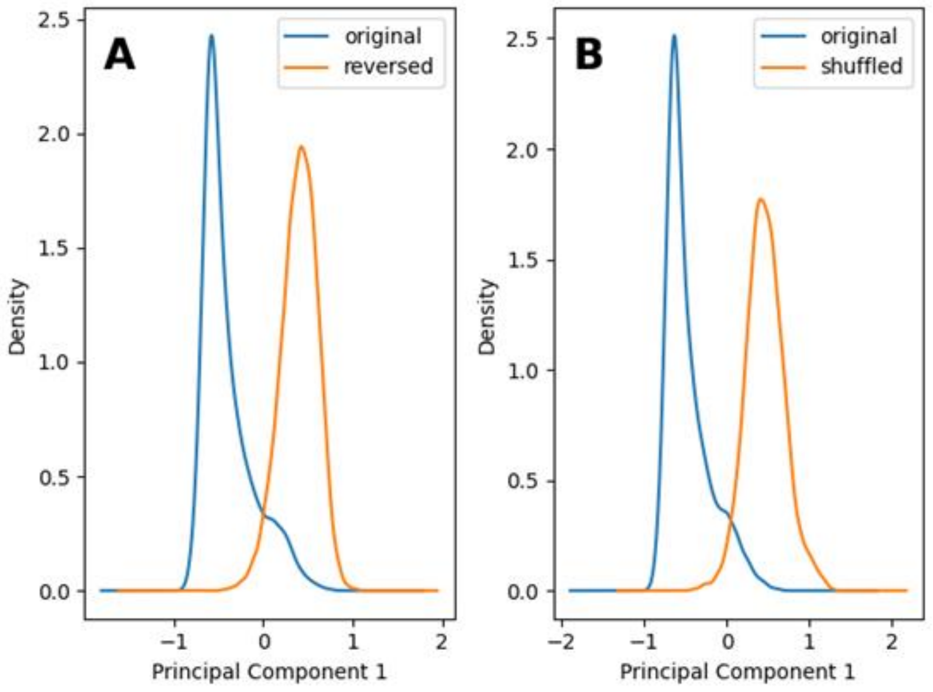

**Fig. S7: Comparing shuffled and reversed sequences.** We apply PCA together to embeddings of original and the same sequences together. We show Kernel Density Estimate of the first Principal Component of the PCA. **A**: original and reversed sequences. **B**: original and shuffled sequences.

**Table S1: Euclidean vs. cosine CATH20 accuracies**

| | *QnormTop1 euclidean* | *QnormTop1 cosine* | *QrawTop1 euclidean* | *QrawTop1 cosine* |
|---|---|---|---|---|
| ProtT5 XL U50 fp16 | 55.3% | 57.5% | 70.1% | 70.9% |
| ProtT5 XL U50 | 55.3% | 57.5% | 70.1% | 70.9% |
| ProtT5 XL U50 L2 | 54.6% | 56.6% | 69.3% | 70.3% |
| ProtT5 BFD | 52.5% | 54.3% | 69.8% | 70.8% |
| ESM1b | 47.7% | 47.9% | 68.3% | 68.5% |
| ESM | 37.9% | 43.5% | 61.6% | 65.2% |
| ProtAlbert BFD | 20.0% | 20.2% | 34.6% | 34.7% |
| SeqVec LSTM1 | 16.7% | 18.6% | 35.1% | 37.4% |
| SeqVec Sum | 16.7% | 18.2% | 35.5% | 37.5% |
| PLUS | 17.5% | 17.7% | 35.7% | 36.0% |
| SeqVec LSTM2 | 16.5% | 17.6% | 35.2% | 36.7% |
| ProtXLNet UniRef100 | 14.1% | 15.4% | 32.3% | 34.2% |
| ProtBert BFD | 11.9% | 12.7% | 20.2% | 21.0% |
| UniRep | 9.1% | 9.1% | 22.3% | 22.4% |
| SeqVec CharCNN | 2.6% | 2.7% | 4.2% | 4.2% |
| AA Composition | 2.5% | 2.5% | 3.9% | 4.0% |
| CPCProt | 1.9% | 2.1% | 3.8% | 3.9% |

*	**Table S1: Euclidean vs. cosine CATH20 accuracies.** Comparing accuracies of different language models for the CATH dataset as explained in Table 1, but also showing the euclidean distance.

**Table S2: Separating fake sequences by PC1**

| | *PC1<0* | *PC1>0* |
|---|---|---|
| *original* | 8856 | 1144 |
| *reversed* | 363 | 9637 |
| | *PC1<0* | *PC1>0* |
| *original* | 9162 | 838 |
| *shuffled* | 219 | 9781 |

* **Table 2: Separating fake sequences by PC1.** We apply PCA to the ProtT5 {Elnaggar, 2021 #66} embeddings of original and the same sequences scrambled together, then count the number of transformed embeddings that are smaller or larger than 0 on Principal Component 1. We do this twice, once original and reversed together and then original and shuffled together, showing the same clear separation for both.
